# Numerous expansions in TRP ion channel diversity highlight widespread evolution of molecular sensors in animal diversification

**DOI:** 10.1101/2021.11.14.466824

**Authors:** Jan Hsiao, Lola Chenxi Deng, Sreekanth Chalasani, Eric Edsinger

**Affiliations:** Salk Institute for Biological Studies; University of California San Diego; University of Edinburgh; Institut Polytechnique de Paris; University of Chicago Marine Biological Laboratory

## Abstract

Transient Potential Receptor (TRP) ion channels are a diverse superfamily of multimodal molecular sensors that respond to a wide variety of stimuli, including mechanical, chemical, and thermal. TRP channels are present in most eukaryotes but best understood in mammalian, worm, and fly genetic models, where they are expressed in diverse cell-types and commonly associated with the nervous system. Here, we characterized TRP superfamily gene and genome evolution to better understand origins and evolution of molecular sensors, brains, and behavior in animals and help advance development of novel genetic technologies, like sonogenetics. We developed a flexible push-button bioinformatic and phylogenomic pipeline, GIGANTIC, that generated genome-based gene and species trees and enabled phylogenetic characterization of challenging remote homologs and distantly-related organisms deep in evolution. We identified complete sets of TRP superfamily ion channels, with over 3000 genes in 22 animal phyla and 70 species having publicly-available sequenced genomes, including 3 unicellular outgroups. We then identified clusters of TRP family members in genomes, evaluated gene models per cluster, and repaired split gene models. We also produced whole-organism PacBio transcriptomes for five species to independently validate our gene model assessment and model repairs. We find that many TRP families exhibited numerous and often extensive expansions in different phyla. Some expansions represent local clusters on respective genomes, a trend that is likely undercounted due to varied quality in genome assemblies and annotations of non-model organisms. Our work expands known TRP diversity across animals, including addition of previously uncharacterized phyla and identification of unrecognized homologs in previously characterized species.

## INTRODUCTION

Organisms innovate and diversify new biologies, systems, and components through evolutionary engineering. Complex biological systems, like the brain, have genetic components that may originate deep in evolution. A clear understanding of their origins, evolution, and diversity can provide new insights and approaches as to their mechanism and function, and can provide a critical foundation to advancing tools in medicine, from developing new disease models to engineering novel genetic technologies. A critical first step is identifying sequence diversity and establishing patterns of gene family evolution across major clades and species of interest, including calls of gene orthology and homology relative to species where functional knowledge is greatest, such as human and genetic models like mouse, fly, and worm in animals. Genome-scale data sets are required for clear phylogenetic resolution and massive growth in sequencing across the Tree of Life now enables broad deep comparative studies that explore genetic diversity but challenges remain, particularly for highly divergent sequences within and between species. In addition, many studies focus on genetic models or a specific clade and/or subfamily of interest that may impart unrecognized artefacts or biases in establishing or interpreting phylogenetic relations and related calls of orthology and homology of sequences in gene families and superfamilies.

The ability to detect and respond to dynamic external features is critical across all scales of an organism and is central to biological function of cells and systems. Sensory transduction systems, which detect and translate these dynamic features enable molecules, cells, systems, and organisms to monitor and respond to their environment. As might be expected given their fundamental nature, sensory transduction systems are central to biology across all domains of life, at scales from molecular to organismal, and similarly utilize primary molecular sensors at the environmental interface. One such superfamily of sensors is the Transient Potential Receptor (TRP) ion channels. These proteins function as multimodal primary and secondary molecular sensors (mechano, thermo, and chemosensing). They are found throughout eukaryotes and form critical components in neural sensory systems and in diverse other cellular and molecular systems in human and animals. Many family members have been studied in human, fly and worm in the context of basic biology and disease. In addition, human TRPA1 and worm TRPN channels have been developed into powerful genetic tools for use in sonogenetics, which uses ultrasound to control mechanosensitive ion channels and downstream physiology and behavior, an emerging technology analogous to optogenetics but without limitations of light and transparency (Ibsen et al. 2015; Duque et al. 2020). However, relationships between families remain contentious, even in human, and diversity within families is inconsistently characterized across animals. This uncertainty adversely impacts design and interpretation of experiments and confounds fundamental questions, like what are the origins of the brain and how have sensory components and systems more generally evolved in organisms? Here, we characterize TRP superfamily diversity in seventy species representing 21 animal phyla and three unicellular outgroups.

## RESULTS

### Published TRP superfamily trees

To assess current knowledge of TRP evolution in animals, seven recently published TRP superfamily phylogenetic trees were characterized for branching structure between families, with selected studies: genome-scale, extending beyond human, fly and worm diversity, and including the classic families TRPA, TRPC, TRPM, TRPML, TRPN, TRPP, and TRPV (Peng et al. 2015; Palovcak et al. 2015; Cattaneo et al. 2016; Kozma et al. 2018, 2020; Kenny et al. 2020). A recent review of the TRP superfamily was also included, as it offers a comprehensive summary of TRP evolution and subjectively considered (based on interpretations of evidence across animals) TRP family branching patterns in a tree schematic, though its actual phylogenetic tree that we also assessed was limited mostly to human, fly, and worms (Himmel & Cox 2020). We find that there is often poor resolution of TRP family relationships in recent studies, with no study providing a fully-resolved tree after collapsing internal nodes based on bootstrap support, that no two studies were in full agreement on inter-family relationships, and that all studies neglected worm Trpml2-Trpml5 and fly TRPL2 TRP genes (Figure 1). We also find that recent comparative studies on TRP evolution and diversity are often imbalanced, with inclusion of just one or a few TRP families or focused on just a major animal clade, plus human (*Homo sapiens*), fly (*Drosophila melanogaster*), and worm (*Caenorhabditis elegans*), and many lack balance across TRPs and/or animals. These results raise the question of why the studies vary in fundamental relationships within the TRP superfamily, highlight the need for new detailed analysis of TRP evolution in animals, and, given the importance of TRPs to understanding human brain and animal biology origins and evolution and potential in sonogenetic technologies, motivated our subsequent work to characterize TRP superfamily gene and genome evolution in animals.

**Figure 1.**
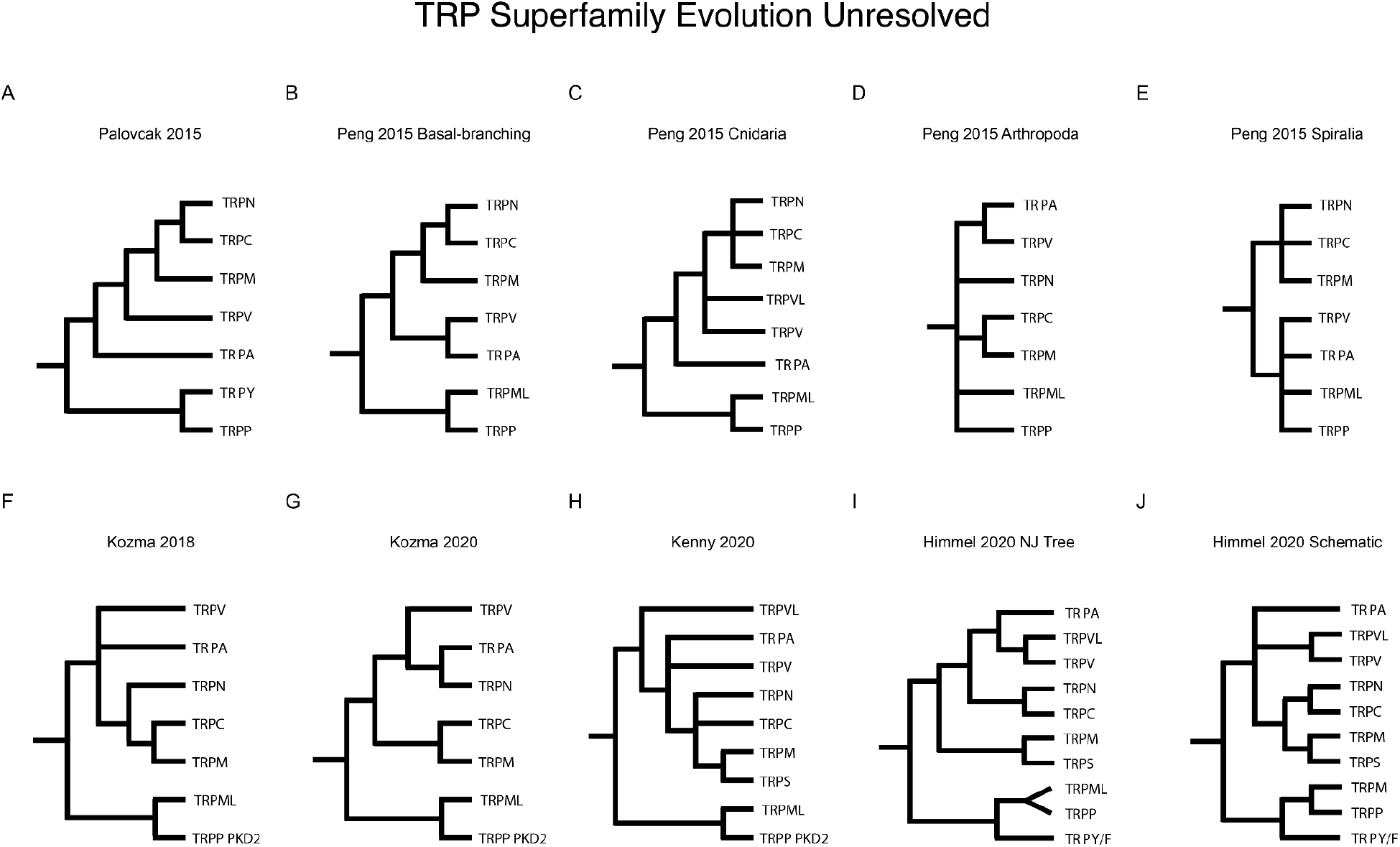
TRP superfamily trees in literature do not fully support one another. **1A**. Palovcak 2015 TRP superfamily tree for animals. **1B-1E**. Peng 2015 TRP superfamily trees for major animal clades. **1F-1G**. Kozma 2018 and Kozma 2020 TRP superfamily trees for arthropod diversity and other animal species. **1H**. Kenny 2020 TRP superfamily tree for sponges and animals. **1I-1J**. Himmel 2020 TRP superfamily trees for animals based on phylogenetic analysis (**1I**) and a schematic summarizing literature interpretation (**1J**).

### Reference Gene Set

We found that human or rodent and distantly-related genetic models fly and worm were common to the seven TRP studies we evaluated and more generally are found in nearly all TRP phylogenetic analyses to date. The human, fly, and worm offer a high-quality minimal representation of animal diversity that includes polished genomes and deep functional knowledge of TRPs. Given these features, we performed initial analyses and developed our informatic pipelines based on a Reference Gene Set (RGS; aka Metazoa3) that included all known TRPs in human, fly, and worm, plus mouse TRP to fill out TRP subfamily diversity in mammals and anemone TRPVL to represent a possible eighth TRP family in animals, for a total of 73 sequences (32 human, 17 fly, 22 worm; 1 mouse, and 1 anemone; Supplemental Material 1). Of note, fly Trpm1 and Trpml2 are identical in DNA sequence and reside at neighboring positions in the genome (coordinates). We included both, though it appears only Trpm1 is used in previous studies and its unclear if Trpml2 is an assembly artefact, pseudogene, or real (see also public database analyses below). Similarly, worm Trpml1-5 are recognized TRP loci in the worm genome, as highlighted in the online WormBook resource but are absent in TRP publications. We included only Trpml2-5 in RGS, as Trpml1 is considered a pseudogene in the genome (see public database analyses below). Overall, we produced a reference gene set that includes some of the most highly-studied TRP sequences in animals and provides a more fully complete representation of TRP diversity in fly and worm genetic models.

### TRP sequence identities in human, fly, and worm

Human TRP sequences are highly divergent between families, with sequence identities as low as 27%. Importantly, sequence divergence is likely a critical factor underlying published differences in family branching structure and TRP superfamily evolution. Sequence alignment methods generally work well given ∼30% or greater sequence identities between sequences but accuracy rapidly declines below this threshold, with standard alignments up to 90% incorrect at 24% identity in comparison to more accurate structural alignments (Haddad et al. 2020). To provide an explicit minimal representation of TRP sequence divergence in animals, we performed an all against all pairwise analysis of RGS TRPs, using water for alignment and calculation of sequence identity. This work produced a total of 5,329 pairwise alignments that included sequence identities ranging from 5% to 100% (Supplemental Table 3). These results highlight the extreme divergence of TRP sequences in human, fly, and worm and suggest that bioinformatic and phylogenetic methods used to characterized TRP superfamily evolution and diversity may benefit from optimization for highly divergent sequences.

### giganticTRP

We next built and optimized bioinformatic and phylogenetic pipelines, based on our inhouse bioinformatic and phylogenomic pipeline, GIGANTIC. Based on this work (detailed below), we established a phylogenomics pipeline, giganticTRP, and used it to characterize TRP superfamily evolution at genome scale and across animals (Metazoa3, Metazoa30, Metazoa63, and Metazoa70). To establish giganticTRP, we focused first on tree building (giganticATP - A Tree Please) and second on homolog identification (giganticOMG - One More Gene).

### giganticATP Metazoa3 RGS optimization for tree building

We surveyed recently published methods and found that a typical TRP family or superfamily tree-building pipeline included: MAFFT sequence alignment, TrimAI alignment trimming, and IQTREE tree building, all with largely default settings. The pipelines were largely standard for the field but notably did not include features or optimizations for deep sequence divergence. Using the typical tools and settings as a reference pipeline, we explored diverse software and parameters using Metazoa3 RGS sequences (Supplemental Table 1; Supplemental Material 1; Supplemental Material 2), including newly available tools and features that had not been previously tested, like the MAFFT -dash option for generation of structural alignment seeds by DASH within MAFFT to improve alignment of divergent sequences (Rozewicki et al. 2019), the ProtSub substitution matrix that is based on deeply divergence sequences (Jia & Jernigan 2021), and novel Clipkit software for alignment trimming and the often strong to superior performance of untrimmed alignments (Steenwyk et al. 2020). We produced hundreds of trees whose family branching structure in the superfamily tree was evaluated (Figure 2) and arrived at the following subset for a second round of more rigorous optimization and pipeline selection: (MAFFT vs MAFFT-DASH alignment) x (full-length vs pore region sequences) x (untrimmed vs TrimAl vs ClipKit -smartgappy alignment trimming) x (Blosum30 vs Blosum45 vs Blosum62 vs Prot2021 substitution matrix) x (IQTREE2 vs FastTree2 tree building). We tested a total of 63 pipelines and their associated trees were collapsed based on bootstrap support and evaluated for family branching structure, similar to before. We found a total of 20 distinct superfamily branching patterns at the family level (Figure 2 3B-3U), with the most highly structured tree (Tree 1 in Figure 2) resulting from a single pipeline: (MAFFT-DASH - pore region sequences - BLOSUM45 substitution matrix sequence alignment) + (ClipKit SmartGap alignment trimming) + (IQTREE2 tree-building). More specifically, for sequence alignment, the top-scoring pipeline used MAFFT-DASH with Dash for initial structural alignment, the BLOSUM45 substitution matrix, and the slow but most accurate linsi settings. For alignment trimming, the new ClipKit tool with the smartgap option worked best for the reference pipeline. Of note, we initially tested IQTREE2 vs FastTree2 using the reference pipeline and found that, similar to a recent study, branching in FastTree2 was often similar to IQTREE2 (though IQTREE2 is superior in resolution based on branch support) but much faster. Thus, for later work in giganticOMG optimization, where tree-building itself was not a focus, we typically used FastTree2 over IQTREE2. Surprisingly, the reference pipeline commonly used in recent studies did not provide the highest resolution of superfamily branching structure. More generally, we find that superfamily branching patterns are sensitive to changes in pipeline tools and parameters and that even the top-performing pipeline fails to fully resolve the tree using a comprehensive collection of human, fly, and worm Metazoa3 RGS TRP sequences. Overall, these results suggest an optimal pipeline for tree-building in giganticTRP but that the deeply divergent nature of TRP sequences across animals remains a challenge, as the pipeline, though optimized for deep divergence and including recent advances in the phylogenetic field, is only slightly better than a number of other pipeline variants tested, offering only a slight gain in tree structure and leaving full-resolution of the tree unachieved.

**Figure 2.**
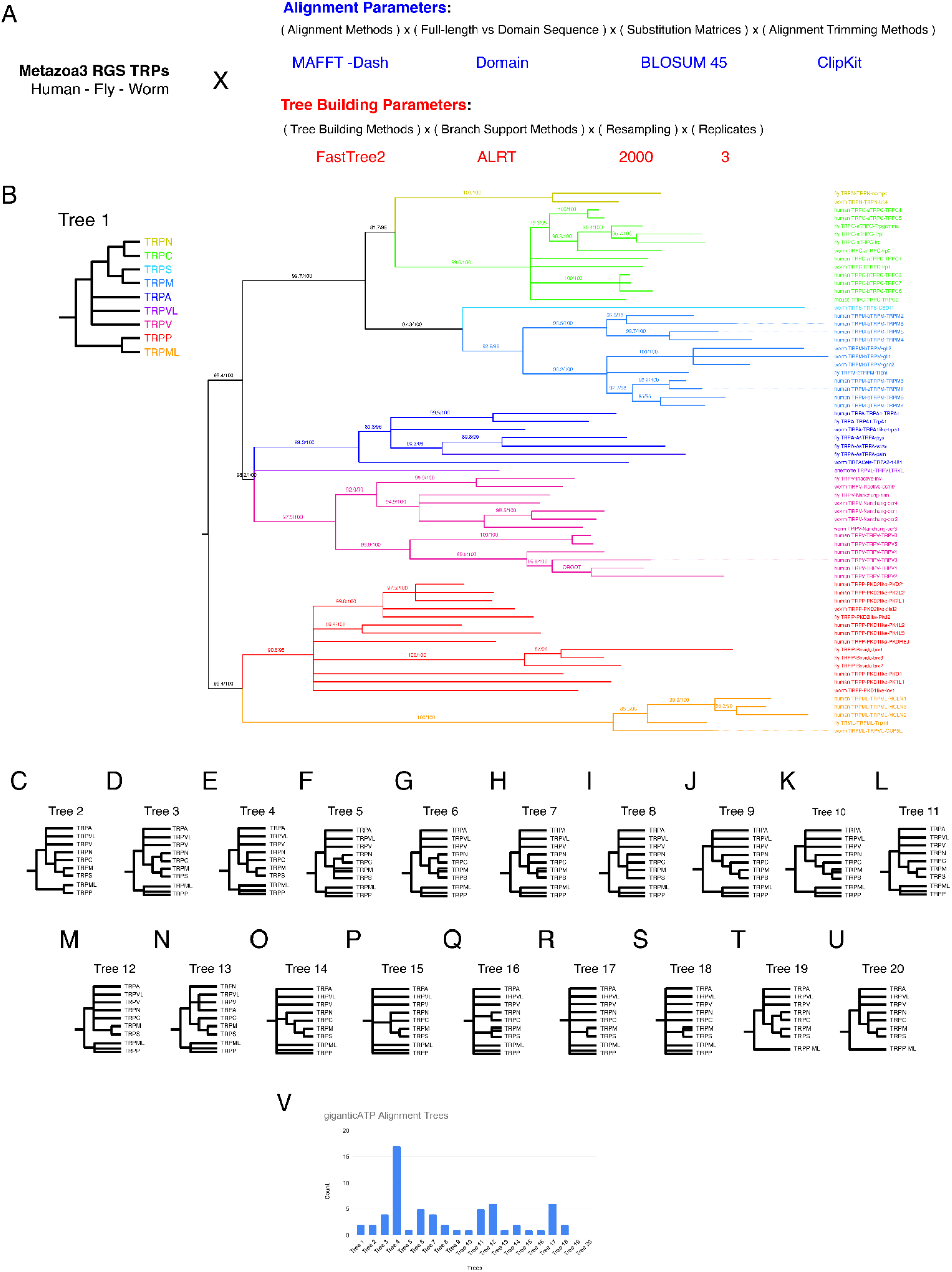
TRP superfamily tree for human, fly, and worm is highly sensitive to alignment and tree-building parameters. **2A**. Alignment and treebuilding parameters that produced the most highly resolved TRP superfamily tree. **2B**. The most highly resolved TRP superfamily tree out of 70 trees produced and with >80% -alrt branch support. **2C-2U**. Additional trees produced for human, fly, and worm TRP sequences when varying alignment and tree-building parameters and with >80% -alrt branch support. **2V**. A histogram summarizing number of trees produced for branching structures represented in 2B-2U with more than 70 trees generated.

**Figure 3.**
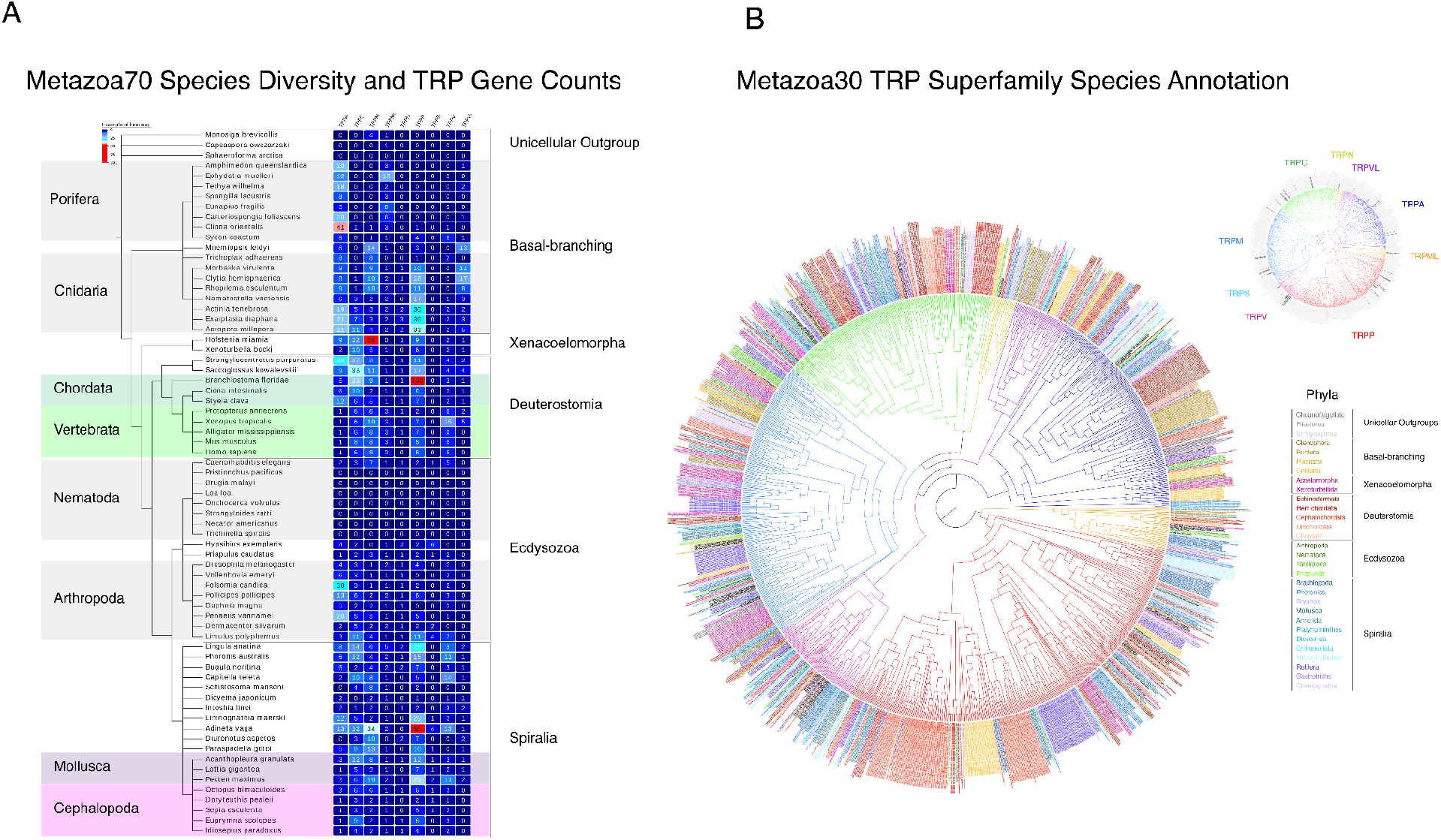
TRP superfamily tree for 21 animal phyla and 70 species. **3A**. Metazoa70 species tree and number of TRP genes identified per family per species. **3B**. Metazoa30 TRP superfamily tree that highlights number gene family expansions per species or phylum.

### Species Tree

To enable interpretations of TRP gene and genome evolution between species and clades, we produced a BUSCO Metazoa-based maximum likelihood species tree for Metazoa70 species and evaluated it in context of ongoing discussions regarding phylogenomic approaches in early animal evolution (Figure 3). BUSCO Metazoa is composed of 972 genes identified as typically single-copy in animal genomes and likely orthologous across species, making them ideal for phylogenetic analysis but largely untested for deep animal evolution. We incorporated recent phylogenetic advances focused on basal-branches in the animal tree and largely untested outside datasets, including use of Clipkit SmartGap for trimming and the LG+C60 sequence evolution model for IQTREE2. We find our species tree is reasonably resolved and largely matches similar studies, though branching at bases of Eumetazoa, Spiralia, and Bivalvia remains ambiguous. An ongoing, if hopefully slowly resolving, controversy that first arose in early phylogenomic work on animal evolution is that ctenophores (*Mnemiopsis leidyi*) vs. sponges (*Amphimedon queenslandica* and *Ephydatia muelleri)* varyingly place as the basal branch in animals and that acoels (*Hofstenia miamia*) varyingly place as the basal branch of Bilateria vs within Deuterostomia between studies. A growing consensus is that placement depends on methodology and tools, and in support of this we find that IQTREE run with LG+C60 places ctenophores at the base of the animal tree and acoels at the base of bilaterian tree. In addition but as expected from previous studies, a number of deep nodes (Ctenophora - All Other Metazoa, Xenocoelomorpha - All Other Bilateria, and base of Spiralia and various nodes within it) have branches that are very short while some species (*Dicyemia japonica, Adineta vaga, Oikopleura dioica, Hofstenia miamia*, and *Mnemiopsis leidyi*, and others) have branches that are very long, suggesting additional genes and alternate species might further improve the tree. Somewhat surprisingly, cephalopods (*Octopus bimaculoides*) place as a sibling branch to gastropods-bivalves (*Lottia gigantea* - *Perna viridis, Mytilus galloprovincialis, Pecten maximus, Saccostrea glomerata, Crassostrea gigas*, and *Crassostrea virginica*), possibly supporting their shared ancestry with monoplacophorans, which were not included. Also of note, coelacanths and lungfish placed on sibling branches to one another (*Latimeria chalumnae* - *Neoceratodus forsteri* and *Protopterus annectens*) and not with coelacanths as sibling to lungfish-amniotes, in contrast to their recent genome projects, though on a very short branch for the node. Unexpectedly, branching between phyla within Spiralia is entirely unresolved and this condition is sensitive, at least in part, to inclusion of nemerteans (*Notospermus geniculatus*, data not shown). Overall, areas of support and ambiguity in our species tree are largely similar to previous studies and the tree provides a phylogenetic framework for interpretation of TRP evolution in Metazoa70 species, albeit with limitations inherent to data and methods currently available.

### TRP families

To characterize TRP gene family diversity and evolution, we produced a series of TRP superfamily trees using the pore region reference gene set and a Mafft-Dash-Clipkit SuperGap-FastTree2 and IQTree2 pipelines (Figure 3 and Figure 4). The Metazoa30 maintained single-species per phylum (Figure 3) and was selected from the broader series of trees that were generated using Metazoa3/30/64/70 gene sets (Figure 4), as it was most useful for visually interpreting patterns of gene loss and expansion in regards to phyla. In contrast increasing species diversity within phyla appeared to provide greater resolution of branching between families for the superfamily tree, as seen in Metazoa64 and Metazoa70 trees versus the Metazoa3 and Metazoa30 trees.

**Figure 4.**
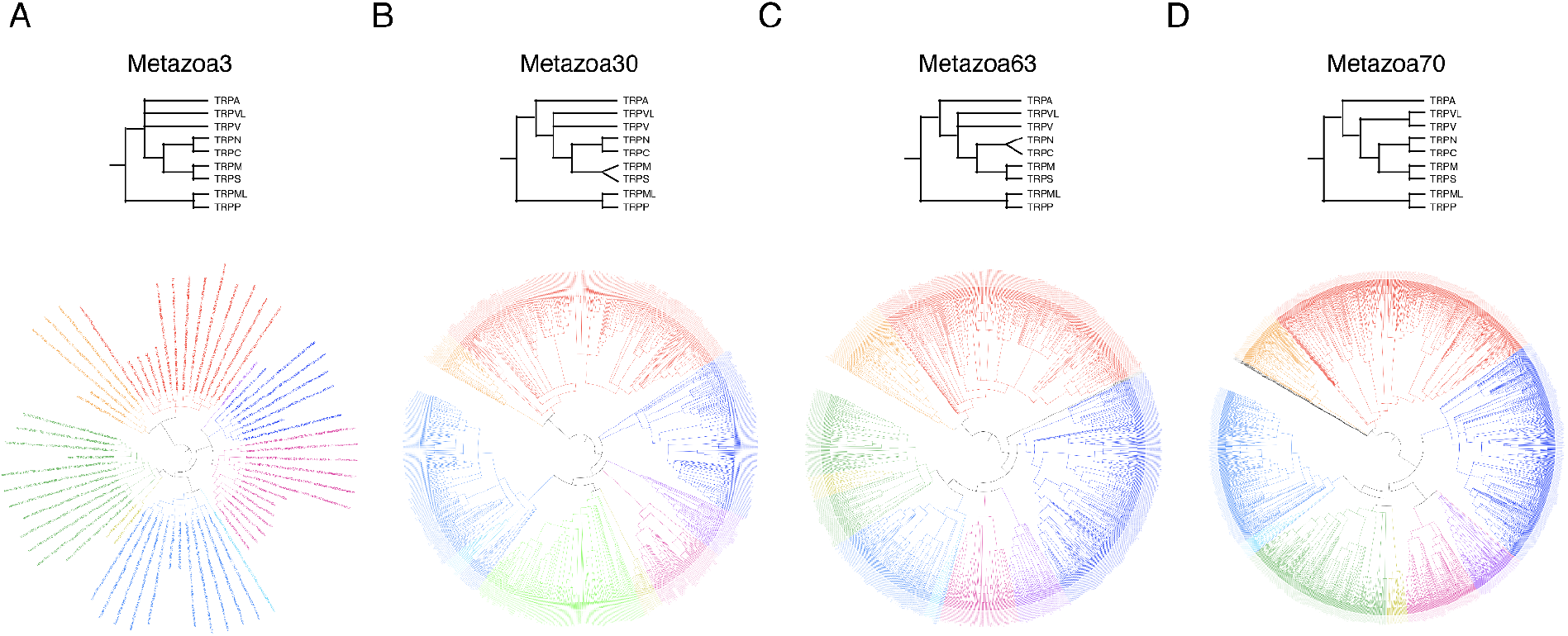
TRP superfamily tree has greater resolution with increasing species diversity but remains sensitive to parameters. **4A**. TRP superfamily tree for Metazoa3 which provides a commonly used minimal representation of animal sequence diversity using human, fly, and worm. **4B**. Metazoa30 TRP superfamily tree. **4C**. Metazoa63 TRP superfamily tree. **4D**. Metazoa70 TRP superfamily tree.

Metazoa70 TRP sequences were assigned to one of eight families (TRPA, TRPC, TRPML, TRPMS, TRPN, TRPP, TRPV, and TRPVL) or a general category of “Unresolved”. To make these assignments, each family was defined on the tree by the deepest node that included all RGS sequences belonging to the family and did not include RGS sequences from another family (Figure 2). Overall, we identified a total of over 3000 sequences that placed in one of the eight TRP families, while a small subset of sequences remained unresolved as to family placement. Unresolved sequences may include novel TRP sequences that are outside traditional families, however, sequences were not highly structured in branching and spot checking revealed that many performed poorly in regards to branch support, potentially due to highly divergent sequence and/or partial sequence. In addition, its likely some subset of sequences maybe false positives that made it through the pipeline without detection. In support of this, we seeded a tree with ankyrin repeat genes from other families and found the false positive sequences placed within TRPs with strong bootstrap support, showing that false positive can place deep within the TRP tree and can be hard to detect based only on tree branching and branch support (data not shown).

As might be predicted, all animal species tested were found to have TRP ion channels, ranging from two families represented in the sponge *Ephydatia muelleri* (TRPA and TRPML) and as few as six channels in the dycemid *Dicyema japonicum* (an unusual clade of parasites known only to live in the enal appendages of cephalopods) to well over half the species having representation of seven or all eight TRP families (FIgure 2). At the same time animals having large complex nervous systems, including independent evolution in cephalopods and vertebrates, have some of the lowest gene counts per family and for all TRPs out of all species examined outside parasitic species. Given the conservative nature of our filtering steps, we expect TRP diversity is underrepresented for many species.

Finally, we assessed more global patterns of TRP diversity per species and per family across animals. Most striking is the common and often extensive occurrence of gene family expansions in diverse species lineages and TRP families. This is visually evident in the Metazoa30 tree, where numerous blocks of color are seen throughout the tree, and each block represents an expansion within the species or its representative clade. We find it is relatively rare that conservation of 1-to-1 orthology is present for a given gene in most animals. Similar to patterns in number of TRPs, cephalopod and vertebrates exhibit among the lowest levels of gene family expansion out of all Metazoa70 species and phyla, a striking contrast to their complex brain and nervous systems.

## DISCUSSION

Overall, we show that TRPs have diversified often and substantially in different families and phyla, regardless and often contrary to nervous system complexity. Thus, while the brain is clearly a sophisticated complex system, as shown here for TRPs, this complexity is not necessarily reflected in the genetic diversity of component gene families in human or octopus, relative to other species. At the same time, organisms lacking similar levels of sophistication in neural tissues and structures often harbor much greater levels of gene diversity for homologous components.

More generally, genome sequencing of diverse organisms has shown that seemingly less complex organisms can have high levels of gene diversity / gene content (Hahn & Wray 2002; Kenny et al. 2020) and this is something we show here clearly and repeatedly across animals for TRPs. In this context, its important to consider that all major groups of living animals represent distinct lineages that, identical to human, have been evolving over the last 550 million plus years since the last common ancestor. In the case of TRPs, they have continued to evolve independently and often massively in different lineages, regardless of and often in contrast to higher levels of organismal structural and behavioral complexity. In this context, the observed complex evolution of TRPs and of potentially other diverse gene families in anatomically or behaviorally simple organisms may reflect cryptic complex biologies that we are only beginning to recognize and understand. Future studies establishing comparative evolutionary frameworks that dissect, compare, and integrate complexity at molecular, cellular, and organismal levels may lead to novel insights and enable a greater appreciation and understanding of diversity and complexity in “simple” vs “sophisticated” systems and organisms.

### TRP Superfamily in animals

TRPs are unusual ion channels in that they are defined strictly based on homology and not by function (Venkatachalam et al. 2014). Failure of our own work to fully resolve phylogenetic relationships of homology and orthology and the lack of consensus in previous studies that we have demonstrated would seem problematic, given the importance of homology in defining TRPs. However, its important to recognized that TRP families themselves are clearly defined with strong bootstrap support (here and in other studies), typically 100% and it is only relationships within and between families that are unable and poorly resolved. Furthermore, many subfamilies, like TRPA1, have strong core branches, though again relationships within and between subfamilies remain poorly resolved and it is possible only a subset of members are included in these core branches, particularly for species that seem prone to rapid sequence evolution and long-branches in the different trees.

Branches in phylogenetic trees are commonly evaluated based on bootstrap support but thresholds for support in traditional methods are arbitrary. Still, above 50% it is generally thought the odds are in favor of a branch. If true, then independent evaluations of branching structure might be expect to converge on a single structure despite low support for some branches. However, our evaluation of TRP superfamily branching structure in the literature indicates that no consensus branching structure exists, even in somes cases within single studies (Peng et al. 2015; Himmel & Cox 2020). Similarly, focusing on human, fly, and worm, where TRP diversity is well characterized, we found a broad diversity of trees are generated for the TRP superfamily when alignment parameters are varied. Importantly, while branching varies substantially between and within families, gene diversity per family is largely unchanged. This result indicates that phylogenetic placement of sequences into families is robust and so current and previous identification of a sequence as belonging to a TRP family is likely correct in most cases, even if relationships within and between families may remain ambiguous. The complexity of expansions and duplications across TRPs and animals, coupled with often poor branch support for various sequences, clades, and/or species, suggests details of sub-family assignment will in some cases be challenging and will require updating as new data and methods advance the field.

### Future Efforts

Molecular sensors, like TRPs, are fundamental to the biology of all living organisms but their role and function outside the nervous system in genetic models is often poorly understood. TRPs are known for multimodal functionality (Venkatachalam et al. 2014) and rapid sequence evolution (Kadowaki 2015)) and this highlights challenges in annotation of TRP diversity based on much more limited functional work in a few species. The numerous massive expansions of TRPs that we have shown here is mind boggling to consider in context of each channel’s specific expression, function, and integration into shared vs novel features of each species and their diverse biologies. Functional and structural work that systematically addresses this diversity is a daunting but likely informative challenge with unexpected payoffs and insights and might be driven through strong collaborative and greater community efforts on emerging new models and large-scale pan-organisms efforts.

## METHODS

### TRP diversity in published studies and public database

#### Published trees: TRP superfamily tree structures

TRP superfamily phylogenetic trees from eight studies were characterized for branching structure at the family level (Peng et al. 2015; Palovcak et al. 2015; Cattaneo et al. 2016; Kozma et al. 2018, 2020; Kenny et al. 2020; Himmel & Cox 2020). Branches were collapsed based on bootstrap support, when available. Threshold for collapse was 95% for ultrafast bootstraps, per guidelines (Minh et al. 2013). Threshold for support is arbitrary in traditional bootstrap methods and found to vary between studies, so branches were collapsed based on each study’s stated cutoff to accurately represent the work as published.

### Public databases: Sponge TRPA1 at Ensembl Compara and NCBI RefSeq

To see how Ensembl characterizes TRPA genes in sponge, Human TRPA1 protein (Uniprot O75762) was blasted online against the sponge genome at Ensembl Metazoa (Release 51 - May 2021) and Ensembl Compara Metazoa gene and family trees for the top hit examined. The same was done for the human genome at Ensembl (Release 104 - May 2021). The sponge genome in Ensembl Metazoa was also keyword searched for ‘TRPA1’.

NCBI SmartBLAST provides broad low-resolution phylogenetic context (alignments and trees) for any sequence using a Landmark Database of 27 high-quality genomes, including human, mouse, fish, fly, and worm in animals, or else NCBI nr. To see how NCBI characterizes TRPA genes in sponge, Human TRPA1 protein (Uniprot O75762) was blasted against the sponge genome in NCBI Genome RefSeq (version 204 - May 2021). The top hit sponge protein sequence was then blasted in SmartBLAST against the NCBI Landmark database and the gene tree examined. The sponge genome in NCBI Genome RefSeq was also keyword searched for ‘TRPA1’.

### GIGANTIC bioinformatic and phylogenetic pipelines

The inhouse GIGANTIC phylogenomic pipeline consists of giganticPDB, which builds a genome databases for the project, giganticOMG, which generates a candidate gene set for species of interest (Metazoa70 in this case), and giganticATP, which produces alignments and gene and species trees. All bioinformatic and phylogenetic pipelines were built using open source software and in-house scripts developed in Unix, Python, and R (Supplemental Table 2). All software was run as default unless noted. GIGANTIC pipeline scripts are available on request and the tool will be published elsewhere.

### giganticID phyloname and phylogenome identifiers

giganticPDB sequence identifiers, giganticIDS, were built based on phylogenetic and source database information. giganticID phyloname identifiers were unique ids for each species. giganticID phylogenome identifiers were unique ids for each genome or transcriptome.

giganticID phyloname ids were structured as:

Kingdom-Phylum-Class-Order-Family-Genus-species_subspecies-NCBI_TaxonId (Metazoa-Chordata-Mammalia-Primates-Hominidae-Homo-sapiens-9606)

giganticID phylogenome ids were structured as:

Phyloname-Source_database_genome_id.sequence_type (Metazoa-Chordata-Mammalia-Primates-Hominidae-Homo-sapiens-9606-GRCh38.aa).

To build identifiers, a standardized, expert-sourced, and up-to-date taxonomy, the NCBI Taxonomy database (https://www.ncbi.nlm.nih.gov/taxonomy)(Schoch et al. 2020), was used (downloaded 14 February 2021). Phyloname identifiers were generated based on the NCBI Taxonomy rankedlineage.dmp file and used to build headers in downstream processing. Phylogenome identifiers were generated based on phylonames and source genome or transcriptome identifiers and used to name fasta files in downstream processing. Maps linking gigaticID identifiers to source identifiers are available on request.

### giganticPDB

Genome databases were built as part of the giganticPDB pipeline and included 1) GenomesDB, a database of high-quality public genomes, 2) OthersDB, a database of additional genome data sets, and 3) ProjectDB, a database of select data sets from GenomesDB and OthersDB and with selection based on user request and/or user-defined thresholds for BUSCO-based genome or transcriptome quality. giganticPDB ProjectDB is used by the giganticOMG pipeline to produce a final All Gene Set (AGS) that consists of user-supplied reference (RGS) and pipeline-identified candidate (CGS) sequences. giganticOMG AGS is in turn used by the giganticATP pipeline in sequence alignment and gene tree-building.

### giganticPDB GenomesDB

The giganticPDB GenomesDB database consisted of publicly-available animal and protist protein gene model data sets in Ensembl and NCBI RefSeq Genomes. All available protein fasta and gff3 files were downloaded on February 13, 2021 for Ensembl 102 (https://uswest.ensembl.org/index.html), Ensembl Metazoa 49 (https://metazoa.ensembl.org/index.html), Ensembl Protists 49 (https://protists.ensembl.org/index.html), and NCBI RefSeq Genomes 204 for Mammalian Vertebrates, Other Vertebrates, and Invertebrates (https://www.ncbi.nlm.nih.gov/refseq/) (Supplemental Table 1). Genome fastas were filtered to retain longest transcript per locus based on gff3 files for ENSEMBL and based on fasta headers for NCBI RefSeq. Filtered data sets were then evaluated by BUSCO using OrthoDB Metazoa10 (metazoa_odb10 2020-09-10 96203a299d818606c8e7e8f74ddf5b59)(Seppey et al. 2019; Zdobnov et al. 2020). In cases where genomes of the same species were present in Ensembl and NCBI, the genome version having a higher BUSCO single-copy complete score was retained, or the NCBI version was used if identical. Finally, filenames were converted to Phylogenome structure and headers were set to a standardized format (>genus-species-gdb_unique_identifier) and formed the GenomesDB database.

### giganticPDB OthersDB

The giganticPDB OthersDB database consisted of specific genomes identified in NCBI Genomes or else where but not included in Ensembl or NCBI Genomes RefSeq (which had been processed as part of GenomesDB), in the literature, and from our own inhouse sequencing and assembly work. Genomes added to OthersDB had either gene models with no isoforms or were easily filtered for longest isoform (T1) based on header information. Genome and transcriptome data sets had filenames converted to phylogenome structure and headers were set to a standardized format (>genus-species-odb_unique_identifier). Processed datasets formed the giganticPDB OthersDB database.

### giganticPDB ProjectDB

A final giganticPDB ProjectDB database consisted of genomes for specific species of interest and previously assessed, filtered, and standardized in production of GenomesDB and OthersDB databases. Specifically, ProjectDB was built by combining GenomeDB and OtherDB datasets filtering for selected species, with species selected based on phylogenetic representation and BUSCO scores. Headers were updated to GIGANTIC identifier format, giganticIDS. A map of GIGANTIC identifiers to source header detail was also generated. BUSCO (Seppey et al. 2019; Zdobnov et al. 2020) was rerun on ProjectDB fastas to link BUSCO analyses to final ProjectDB identifiers. Blast (Buchfink et al. 2015) databases were built for peptide gene sets of each ProjectDB species. Finally, five ProjectDB species diversity sets were established and consisted of 3,15, 30, 48, and 70 species: Metazoa3, Metazoa30, Metazoa63, and Metazoa70.

### Reference Gene Sets

A TRP superfamily reference gene set (RGS) of 73 proteins was collected based on human, fly, and worm TRP genes, as identified in Kozma 2018 and Himmel 2020, plus representative anemone (TRPVL) and mouse (TRPC2) sequences identified in Himmel 2020(Kozma et al. 2018; Himmel & Cox 2020), and supplemented with known but missing fly (Trpl2) and worm sequences (Trpml2-Trpml5). Actual sequences were from Uniprot (https://www.uniprot.org/; February 2021), identified by searching gene and species names (Supplemental Material 1). Headers in the RGS fasta file were updated to RGS structure: >name-TRP_family-TRP_subfamily-sequence_name-identifier (>human-TRPA-TRPA1-TRPA1-uniprotXXXXXX). The pore region is required for any protein having ion channel function and is both characteristic and conserved in TRPs, while other features are not found in all families and some, like ankyrin repeats, are present in a number of unrelated families. Thus, we also produced a reference gene set fasta of just the pore region per sequence (Supplemental Material 2). To do this, human TRPA1 was graphically annotated based on annotation coordinates, including transmembrane domains (TM) and pore channel, provided in Uniprot based on its structural study (Paulsen et al. 2015). The annotated sequence was aligned with all other RGS TRP sequences by MAFFT in Geneious Prime and a pore region was defined as sequence spanning the six characteristic TM domains, plus highly conserved flanking regions, identified as plus 40 amino acids 5’ of the first TM domain and 40 amino acids 3’ of the sixth TM domain. The remaining 66 RGS TRP proteins were graphically annotated for TM domains in Geneious and the pore region sequence extracted per protein to produce the TRP superfamily extracted pore region reference gene set (RGS-X) (Supplemental Material 2). Although six TMs are expected for TRP channels (or 11 for TRPP PKD1-related channels), predicted TMs ranged from 4 to 13 and boundaries for the pore region were subjectively judged based on alignment and known features of a given family, when necessary. Himmel 2020 (Kozma et al. 2018; Himmel & Cox 2020) initial reference gene set was later found to be missing five worm TRP sequences. These sequences were added and formed reference gene set “RGS TRP”

### Sequence Identity

Sequence identities were calculated based on all against all pairwise alignments of RGS TRP and of RGS TRP X sequences using water as implemented in SeqDiva and visualized using SeqDiva(Agüero-Chapin et al. 2019) and in Geneious.

### giganticOMG

Candidate genes were identified in Metazoa70 genomes using Blastp-based and HMMer3-based methods. Blastp pipelines initially searched RGS73 protein sequences against Metazoa70 genomes. All hits were then blasted back against the reference genomes (human, fly, worm, mouse, and anemone), and all hits having one of the RGS sequences as a top hit were retained. Blastp pipelines varied in Blosum matrix (30, 45, or 62) and use of full-length or pore region RGS73 sequences. A novel HMMer3 Jackhmmer pipeline was also built to identify highly divergent sequences, given the greater sensitivity of HMM-based methods. Pore regions of RGS sequences were used to seed an initial round of Jackhmmer across the Metazoa70 genomes. If the first hit in a reference genome was not an RGS sequence, the hit was dropped and not included in RGS. All new hits with an evalue less than the last RGS sequence were retained and evaluated in a subsequent round of Jackhmmer. Iterations continued until no new sequences were discovered and a final CGS then produced. CGS from Blast and HMMer pipelines were combined, made unique to form the final CGS gene set. RGS73 sequences were added to final CGS sequences to produce a final AGS dataset that was then used in optimized giganticATP pipelines for tree-building.

### giganticATP

#### Published tree-building methods

An initial alignment and tree-building pipeline, giganticATP Baseline, was based on recent TRP papers, with parameters commonly used in gene family phylogenetic studies, and included: MAFFT(Katoh & Standley 2013) alignment, TrimAl (Capella-Gutiérrez et al. 2009) trimming, and IQ-Tree2 (Minh et al. 2020) maximum likelihood tree building. The pipeline provided a reference and platform for optimization of alignment and tree-building (see below).

### giganticATP alignment optimization

To optimize the giganticATP Baseline tree-building pipeline, we initially explored sequence alignment parameters. In particular, given low sequence identity between TRP families (see Results), sensitivity of alignment to low-sequence identity, and sensitivity of trees to alignment, and because alignment has not been explicitly examined in previous TRP studies, we explored sequence alignment parameters, using giganticATP Baseline and RGS TRP and RGS TRP-X reference gene sets. We tested 1) full-length vs. pore region sequences, 2) sequence-based aligner MAFFT vs. sequence-structure-based aligners Promals3D (http://prodata.swmed.edu/promals3d/promals3d.php) (Pei & Grishin 2014) and MAFFT-Dash (https://mafft.cbrc.jp/alignment/server/) (Rozewicki et al. 2019) and also the structure-only aligner mTM-align (https://yanglab.nankai.edu.cn/mTM-align/) (Dong et al. 2018), which was used to provide a constraint tree to Promals3D, 3) amino acid substitution matrices, including BLOSUM30, BLOSUM45, and BLOSUM62, and structure-based ProtSub (Jia & Jernigan 2021), and 4) alignment trimming as untrimmed, TrimAl(Capella-Gutiérrez et al. 2009) trimmed, and ClipKit (Steenwyk et al. 2020) trimmed. Structural data sets for mTM-align, Promals3D, and MAFFT-Dash were selected by searching RCSB PDB (https://www.rcsb.org/) (Berman et al. 2000) for family name (except TRPA1 was used for TRPA) and selecting the dataset with the top structural score when more than one was available. No structures were available for TRPVL and TRPS (PDB searched on 8 April 2021). Promals3D (Pei & Grishin 2014) was run with and without a constraint tree produced by mTM-Align (Dong et al. 2018). MAFFT-Dash (Rozewicki et al. 2019) was run standard, standard with “Homologs” setting and UniRef database, and standard with “Homologs” and “Leave Gappy” settings, and was limited to just pore region sequences, as it requires global homology across sequences. All three structural aligners were run online. Trees were build using IQ-TREE2 with 2000 resamples for ultrafast bootstraps and -alrt. Each tree was evaluated online at the interactive Tree of Life (https://itol.embl.de/) (Letunic & Bork 2019), ultrafast bootstraps less than 95% were collapsed, and each family was color annotated in iTOL or in FigTree.

Two distinct alignment approaches used in previous TRP superfamily gene trees were tested for the Metazoa70 gene set: 1) sequence-based alignment (Peng et al. 2015; Kozma et al. 2018, 2020; Himmel & Cox 2020) and 2) sequence-structure-based alignment (Kenny et al. 2020). For sequence alignment, we used standard MAFFT. For sequence-structure alignment, we used MAFFT -Dash-beta, which enabled structural alignment, outperformed ProMals3D in our hands for Metazoa3, and accommodated our large number of sequences (∼3,000).

### giganticATP tree-building optimization

To initially optimize tree-building, we focused on resampling for ultrafast bootstrap and -alrt branch assessments. We optimized resampling using one of the parameter sets that generated the most highly structure superfamily tree (Tree 1; Pore region x Blosum45 x ClipKit SmartGappy x Maftt -Dash) and performed 2,000, 5,000, 10,000, 15,000, 20,000, and 30,000 rounds of resampling and 10 separate runs for each level of resampling and branch support methods as UFBS, ALRT, or both UFBS and ALRT. Trees were assessed and branches having less than 95% ultrafast bootstrap or 80% ALRT support were collapsed. Trees were then characterized for superfamily branching and evaluated for convergence between replicates per resampling level in iTOL to identify resampling threshold for tree converge in replicates. Based on our optimized resampling threshold, we reran the 63 alignment parameter trees and characterized superfamily tree branching a second time to reevaluate alignment optimization under optimized resampling.

Tree building software using ML methods (IQ-TREE, RAxML, PhyML) outperform those using approximately ML methods (FastTree2) in various metrics (Zhou et al. 2018). However, FastTree2 is up to 1000x faster and its tree topology is highly similar to ML trees for all ML branches having strong support but do suffer in branch lengths (Price et al. 2010; Young et al. 2020). Still, subsequent correction of branch lengths using ML methods produces trees that are of equal quality to a standard ML pipeline (Young et al. 2020). Thus, we explored use of FastTree2 to search tree space and generate an initial tree whose topology was used to constrain IQ-TREE2 (-g setting), which was then used to correct branch lengths. Branch support of ML IQ-TREE2 trees was evaluated both for ultrafast bootstraps of (95% threshold) and SH-aLRT (80% threshold), as the two approaches complement one another, per methods and software guidelines (Guindon et al. 2010; Minh et al. 2013, 2020) (see also IQ-TREE2 documentation: http://www.iqtree.org/doc/). FastTree2 uses approximately ML methods and offers Shimodaira-Hasegawa test to evaluate branch support, with values that correspond to SH-like methods of aLRT SH-like methods in PhyML3, which is in turn shown to be similar to SH-aLRT in IQ-TREE (Price et al. 2010; Guindon et al. 2010; Minh et al. 2020) (see also online documentation: http://www.microbesonline.org/fasttree/). Based on these similarities, we used a similar cutoff threshold of 80% SH value for branch support. FastTree2 SH branch values were transferred to the IQ-TREE2 tree prior to visualization.

### Metazoa70 Species Trees

BUSCO Metazoa-based species trees were generated for the Metazoa63 species diversity set and following recent guidelines (Steenwyk et al. 2020; Pandey & Braun 2021). BUSCO Metazoa genes were aligned all species per gene using MAFFT (Katoh & Standley 2013). Alignments were trimmed using ClipKit (Steenwyk et al. 2020) (default settings, plus Smart Gappy). Sequence evolution models were calculated per BUSCO Metazoa gene alignment using IQ-TREE2 ModelFinder (Kalyaanamoorthy et al. 2017; Minh et al. 2020). Alignments were then concatenated to produce a supermatrix and used to generate a ML supermatrix species tree (Minh et al. 2013, 2020; Chernomor et al. 2016; Kalyaanamoorthy et al. 2017).

### Metazoa70 Gene Trees

Contrast-based color palettes for species and gene trees were per Metazoa70 species and per TRP family, with families including TRPA, TRPM, TRPML, TRPN, TRPP, TRPS, TRPV, and TRPVL.

## LEGENDS

**Supplemental Table 1. Genome and genome publication source details per species used to generate the Metazoa70 and other species gene sets for TRPs**.

**Supplemental Table 2. Software sources used in the GIGANTIC pipeline**.

**Supplemental Table 3. Sequences identifies calculated for all RGS73 sequences in human, fly, and worm**. Sequence identities were calculated by pairwise alignment in water.

**Supplemental Material 1. Reference Gene Set 73 full-length fasta file**. A fasta file of reference gene set full-length sequences for RGS73. Sequences were collected from Uniprot.

**Supplemental Material 2. Reference Gene Set 73 pore region fasta file**. A fasta file of reference gene set pore region sequences for RGS73. Sequences were collected from Uniprot from pore regions of RGS73 pore regions as identified by alignment.

